# Targeted mutations in IFNα2 improve its antiviral activity against various viruses

**DOI:** 10.1101/2023.09.06.556557

**Authors:** Zehra Karakoese, Vu-Thuy Khanh Le-Trilling, Jonas Schuhenn, Sandra Francois, Mengji Lu, Jia Liu, Mirko Trilling, Daniel Hoffmann, Ulf Dittmer, Kathrin Sutter

## Abstract

During viral infections, type I interferons (IFN) are induced and play a key role in counteracting initial viral spread. Twelve different human IFNα subtypes exist that bind the same receptor; however, they elicit unique host responses and display distinct potencies of antiviral activities. Our previous studies on HIV and HBV demonstrated that the clinically used IFNα2 is not the most effective one among the IFNα subtypes. By sequence modeling, we identified a region in helix B with mainly conserved residues at the outside facing IFNAR1, but variable residues at the inside facing the core of IFNα, potentially representing a putative tunable anchor to tune pleiotropic IFN responses. Using site-directed mutagenesis various mutations were introduced into the IFNα2b backbone targeting sites which are important for binding to IFNAR1 and IFNAR2, the putative tunable anchor, or outside these three regions. Selected mutations were based on sequence differences to high antiviral subtypes IFNα6 and IFNα14. Treatment assays against HBV and HIV identified several critical residues for the antiviral activity of IFNα mainly in the IFNAR1 binding region. Combined mutations of the IFNα2 IFNAR1/2 binding sites or the IFNAR1 binding region plus the putative tunable anchor by those of IFNα14 further augmented activation of different downstream signaling cascades providing a molecular correlate for the enhanced antiviral activity. We describe here important functional residues within IFNα subtype molecules, which enabled us to design novel and innovative drugs that may have the potential to be used in clinical trials against a variety of different viral infections.

**Importance:** The potency of IFNα to restrict viruses was already discovered in 1957. However, until today only IFNα2 out of the 12 distinct human IFNα subtypes has been therapeutically used against chronic viral infections. There is convincing evidence that other IFNα subtypes are far more efficient than IFNα2 against many viruses. In order to identify critical antiviral residues within the IFNα subtype sequence, we designed hybrid molecules based on the IFNα2 backbone with individual sequence motifs from the more potent subtypes IFNα6 and IFNα14. In different antiviral assays with HIV or HBV, residues binding to IFNAR1 as well as combinations of residues in the IFNAR1 binding region, the putative tunable anchor, and residues outside these regions were identified to be crucial for the antiviral activity of IFNα. Thus, we designed artificial IFNα molecules, based on the clinically approved IFNα2 backbone, but with highly improved antiviral activity against several viruses.

## Introduction

The importance of interferons (IFNs) in fighting viruses was first discovered in 1957 by Isaacs and Lindenmann showing the potential of IFNs to inhibit virus replication (1). Later, it became apparent that the biological properties of IFNs are much more comprehensive than only their antiviral activity including antiproliferative and immunomodulatory activities. The interferons are clustered into three different types based on sequence homology, receptor usage, and downstream signaling cascades. Even though all IFNs play a crucial role in the fight against viral infections, only type I IFNs are clinically approved and are therapeutically applied to treat chronic viral infections, like Hepatitis B Virus (HBV) and at least in the past Hepatitis C Virus (HCV) (2, 3). Type I IFNs can be affiliated to a multigene cytokine family encoding numerous genes for IFNα, but only a single gene for IFNβ, IFNε, IFNκ, and IFNω. The overall 13 human IFNα genes express 12 different proteins (subtypes) which share a highly conserved protein sequence with up to 95% identity (4, 5). However, they differ markedly in their biological activities (6–11). Interestingly, all IFNα subtypes evoke their full biological spectrum through binding to the same receptor complex, composed of the two transmembrane proteins IFNAR1 and IFNAR2. Downstream signaling is initiated by IFN binding to the high-affinity subunit IFNAR2, which further recruits IFNAR1 to form a ternary complex (12, 13). Different binding affinities to both receptor subunits, as well as the membrane surface concentration of the receptor subunits, and the equilibrium between binary (IFNAR2-IFN) and ternary (IFNAR2-IFN-IFNAR1) complexes on the surface determine the diverse biological response of the different subtypes (12–14). In addition, the diversity of the type I IFN-mediated responses might be further modulated by the type of cell, the microenvironment and the timing relative to other stimuli, e.g., T cell receptor triggering and basal expression of signal transducers and activators of transcription (STAT) (15, 16). The ligation of the ternary complex leads to the activation of the classical Janus kinase (JAK)-STAT pathway which induces the transcription of hundreds of IFN-stimulated genes (ISGs). However, several other pathways, including the mitogen activated protein kinase (MAPK) pathway, the phosphoinositide 3-kinase (PI3K) pathway, and the NF-kB cascade are also activated upon IFN binding, which further tune the pleiotropic IFN response (17). However, the interplay of the IFNα subtype binding, the complex downstream signaling events, and the subsequent biological activities has not been sufficiently investigated yet.

Type I IFNs can be classified as helical cytokines. Their structure consists of five α-helices -from the N-terminus helix A to the C-terminus helix E- which are associated by an overhand loop (AB loop) and three short segment loops (BC, CD, and DE loops) (18, 19). The core of type I IFNs including all helices as well as parts of the AB loop is defined by conserved structures. However, major structural differences between the IFNα subtypes in the AB loop, helix B, and BC, which might account for their different biological activities.

In previous studies, we could show that IFNα6 and IFNα14 had superior antiviral activity against HBV and HIV compared to IFNα2 (9, 11, 20). Especially, IFNα14 has predominant immunomodulatory capacities against HIV by activating effector T and NK cell functions *in vitro* and *in vivo* (7, 11). Further during HBV infection, IFNα14 can activate both type I and type II IFN signaling pathways, resulting in the expression of an enlarged pattern of IFN gamma-activated sites (GAS)- and IFN-stimulated response element (ISRE)-driven ISGs (9) which define the biological activity.

To improve the antiviral activity of IFNα2b we identified critical residues within the IFNα subtypes which might be potentially important for an augmented antiviral activity. Using site-directed mutagenesis various amino acids from IFNα6 or IFNα14 were introduced into the IFNα2b backbone. Antiviral assays against HBV and HIV could manifest the importance of certain regions within the IFNα structure. Further, IFNα2-mutants increased the activation of various downstream signaling pathways making these critical residues potential target regions to improve IFN-mediated responses.

## Results

### Human IFNα subtypes: sequences and structure

The type I IFN family is composed of multiple IFNα subtypes with non-redundant biological activities. So far, only IFNα2a/b was approved for clinical treatment; however, with narrow treatment options and a variety of undesirable side effects (21).

To identify critical residues within the IFNα subtypes which might be potentially important for clinical application, we aligned all IFNα subtypes with multiple alignment using fast Fourier transform (MAFFT) (22). Thereby we aimed to identify potential mutable motifs within the protein structure which could be modified to improve the effector function of IFNα2. The alignment of all human IFNα variants showed in 190 total positions 93 identical positions (’*’; 49%), 46 conserved substitutions (’:’), and 12 semi-conserved substitutions (’.’), resulting in a high total homology of 79%. The root-mean-square deviations (RMSDs) between tested 3D structures of different IFNα subtypes (here IFNα1/13 and IFNα2) were around 1Å. The smallness of deviations between 3D structures (low RMSDs) was not surprising, given the high sequence homology. To see whether the sequence deviations were associated with 3D structural positions, we mapped the sequence entropies computed for the multiple sequence alignment onto the IFNα2 structure (PDB entry 1itf, selected model 1, (19)), (Fig. 1A; Supp. Table 1). The highest entropies (orange and red) were associated with positions that were solvent exposed. The lowest entropies (deep blue) were at positions that were dispersed throughout the structure, with a few remarkable clusters, namely the two C-terminal helices. Interestingly, not only buried positions had low entropies but also exposed positions, probably conserved anchor position involved in receptor binding. Conversely, not all higher entropy positions were exposed, but some were buried, e.g., T14 or F151, possibly tuning the overall IFNα structure.

**Figure 1:**
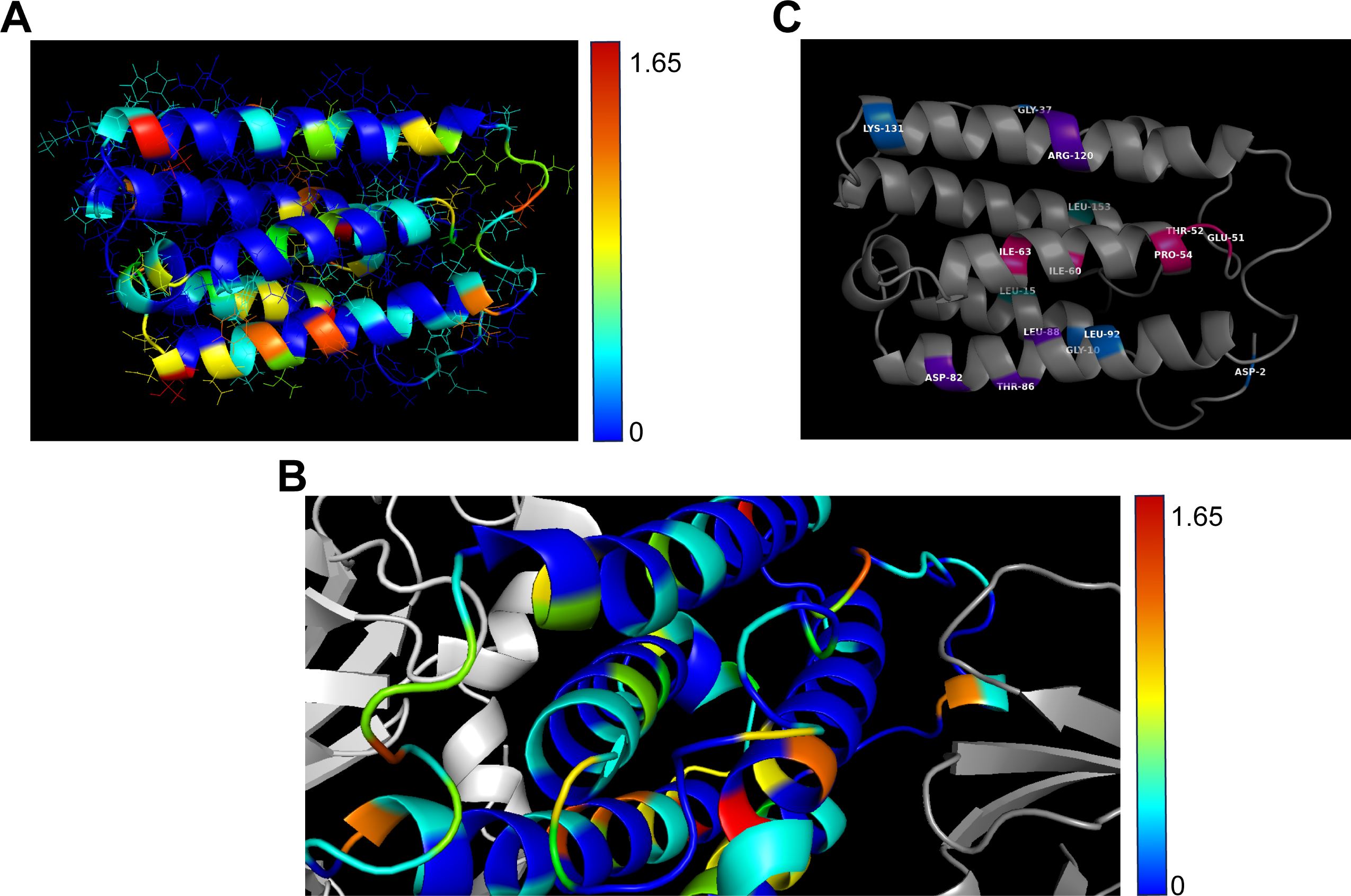
Structural analysis of IFNα subtypes. (A) Entropies encoded as rainbow colors from 0 (blue, complete conservation) to 1.65 (red, highly variable) were mapped on experimental structure of IFNα2 (19). (B) Close-up of ternary complex of IFNAR1 (left, light grey), IFNα2 (middle, after optimal superposition with IFNα2), IFNAR2 (right, dark grey). Colors on IFNα2 mark entropy as in (A) based on PDB entry 3se3 (13). Right of the center lies the conserved anchor helix C138-S150 (blue) that bridges IFNα and IFNAR2 (dark grey). The helix left of the center is the putative tunable anchor with the conserved (blue) outside facing IFNAR1 (light grey) and the variable (green) side facing the core of IFNα. (C) Structure of IFNα2 with key amino acids highlighted in purple (IFNAR1), teal (IFNAR2), pink (putative tunable anchor), or blue (outside).

**Table 1.**
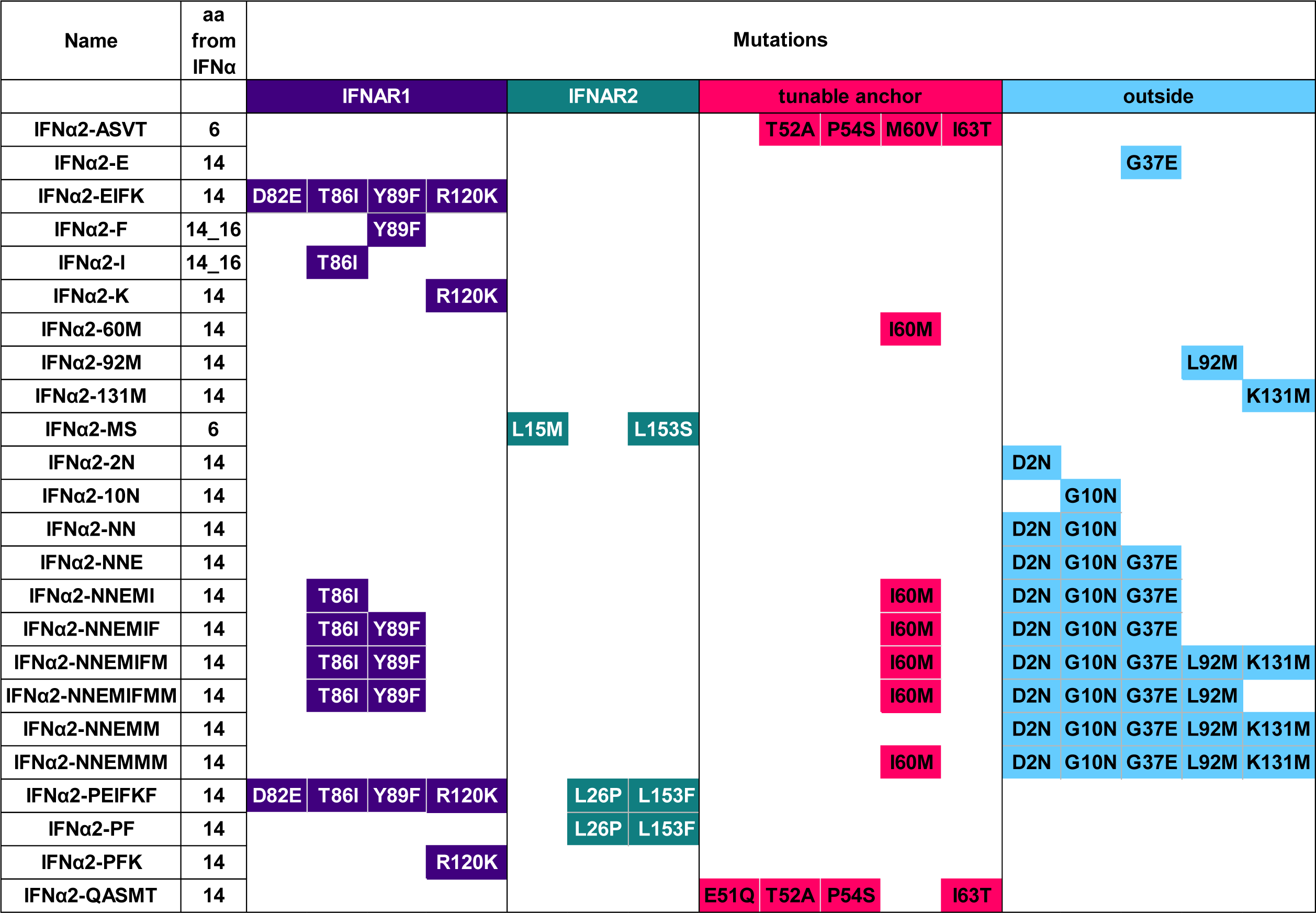
List of IFNα2-mutants.

We also mapped the IFNα1/13 structure with entropies onto the IFNα2 structure in the ternary complex with a partial IFNAR1 and IFNAR2 complex (RMSD between IFNα1/13 and IFNα2: 1.1Å) to address if especially conserved parts of the IFNα were in contact with the two receptor subunits (Fig. 1B). We did not observe a simple relation between parts of IFNα in contact with the receptors and sequence entropy: contacts can be formed by conserved parts (blue) or by variable parts (yellow, orange, or red). Two remarkable helices were identified. First, the helical 4 region C138-S150 close to the C-terminus (Fig. 1B), which is completely conserved in all human IFNα subtypes, and it bridges IFNAR2 with the core of the IFNα molecule. This could be a conserved “anchor”. Second, there was a helix, about T52-L66, with a surprising conservation pattern, namely conserved residues (blue) at the outside but more variable (cyan) at the side facing the core of IFNα. This is a putative “tunable anchor” (TA), i.e., while the binding site with IFNAR1 is conserved, its position and fine structure can be tuned by mutations in the core. Mutations in the putative tunable anchor region of IFNs might positively affect the binding affinity to the IFNAR1/2 receptor and thereby improve their biological activity.

### Targeted mutations in IFNα2 to augment antiviral activity

From the multiple sequence alignment and the analysis of the 3D structure of IFNα within the ternary complex, we identified conserved and variable positions in the receptor binding domains of IFNα as well as a putative tunable anchor which might change the fine structure of IFNα by single or combined amino acid changes (highlighted in Fig. 1C). This might have functional consequences for the IFNα subtypes, especially for their antiviral potential. Since we observed amino acid differences in these regions between IFNα subtypes and previously described differences in their antiviral activities (9, 11, 20), we addressed the question if amino acid exchanges in these regions between subtypes influence antiviral properties. Thus, we produced a variety of different, not naturally occurring IFNα2-mutants, which were all based on the IFNα2b sequence, with specific single or multiple mutations in either the receptor binding site to IFNAR1 (shown in purple), to IFNAR2 (shown in teal), the putative tunable anchor (shown in pink), or outside of these defined regions “outside” (shown in blue) (Tab. 1; Supp. Fig. 1). Since IFNα6 and IFNα14 were the most potent antiviral subtypes in our previous studies (6, 9, 11, 20), we replaced the IFNα2b sequence in the regions mentioned above using site-directed mutagenesis with that from these two potent subtypes, with the goal to augment IFNα2 activity.

### Modified IFNAR binding sites increased antiviral activity of IFNα2 against HBV

We could previously show that IFNα6 and IFNα14 had superior antiviral activity against HBV compared to IFNα2a (9). IFNα14 concurrently activates the type I and type II IFN signaling pathways resulting in the expression of an enlarged pattern of GAS- and ISRE-driven ISGs (9). To define which specific motifs of the protein influence its antiviral activity against HBV, we infected fully differentiated HepaRG cells with HBV and treated the cells with different parental IFNs (IFNα2b, IFNα6, or IFNα14) or the described mutants. In line with our previous results, we observed a strong reduction in HBsAg levels after treatment with IFNα6, whereas IFNα2b had only a minor effect on HBV replication (Fig. 2A, B). Amino acids that are important for binding to IFNAR1 are identical between IFNα2b and IFNα6; thus, only IFNα2-mutants carrying the amino acids of IFNα6 at the IFNAR2 binding site or the putative tunable anchor were analyzed. IFNα2- MS (with IFNAR2 binding sites from IFNα6) and IFNα2-ASVT (mutated in the putative TA region) had no improved anti-HBV activity compared to the parental IFNα2 indicating that modulating the IFNAR2 binding site or the putative tunable anchor alone was not sufficient to enhance the antiviral properties of IFNα2 to the potency of IFNα6.

**Figure 2:**
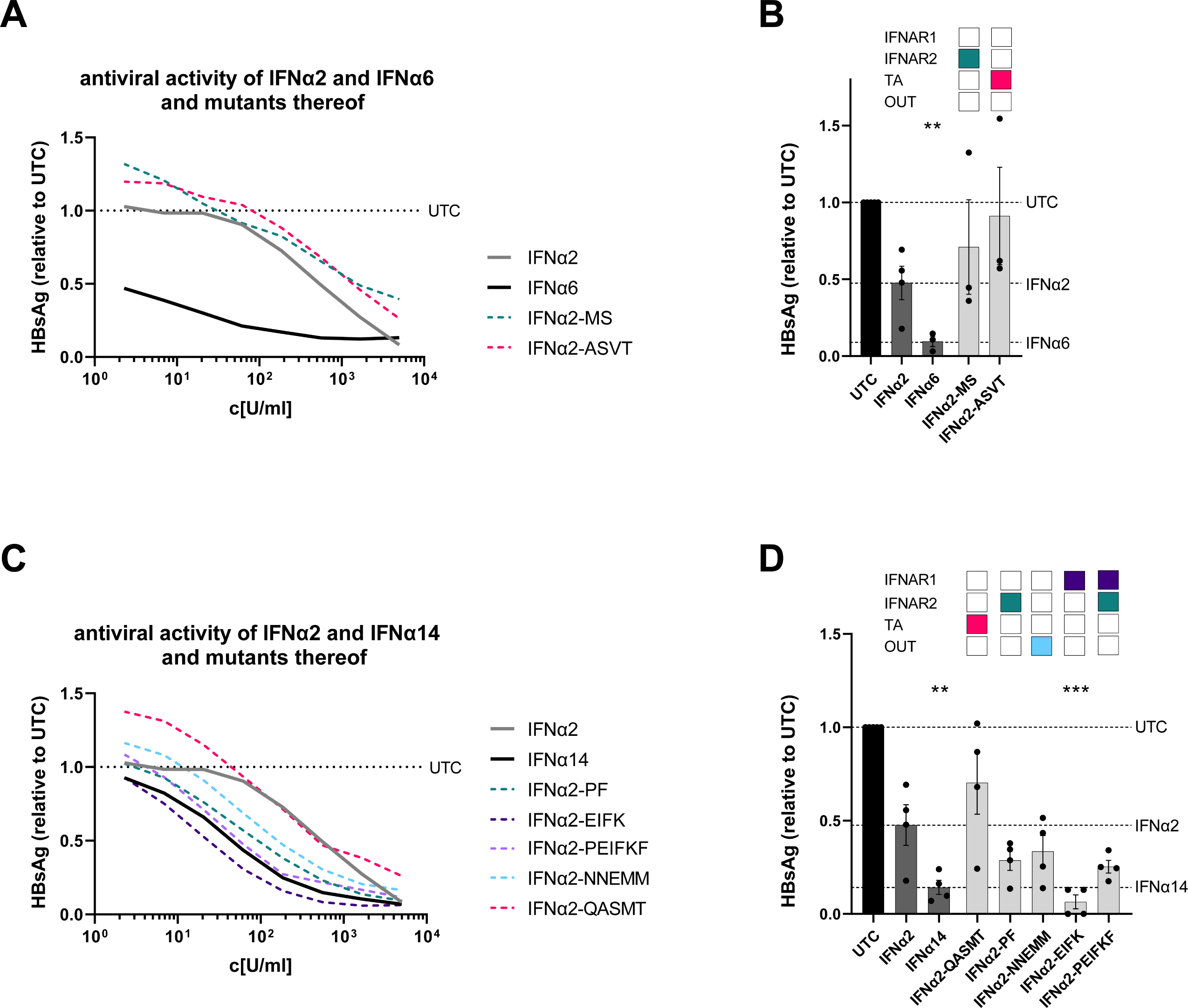
The antiviral activity of IFNα2-mutants against HBV. Differentiated HepaRG cells were infected with HBV at a MOI of 500 geq and treated with different concentrations of IFNα2, IFNα6, IFNα14, or IFNα2-mutants. (A, C) IFNα2, IFNα6, IFNα14, and IFNα2-mutants were serially diluted to measure dose-dependent anti-HBV effects after 8 days by quantifying HBsAg. (B, D) dHepaRG were pre-treated with 2,000 U/ml of the designated IFNs and IFNα2- mutants to perform a screening for their anti-HBV activity by quantifying the amount of HBsAg after 4 days. (A, C) n = 3; (B, D) n = 4; Mean ± SEM is plotted. Kruskal-Wallis test was performed Statistical significance is depicted as ****p<0.0001, ***p<0.001, and **p<0.01. TA = Putative tunable anchor; OUT = outside

Next, we tested IFNα2-mutants harboring distinct amino acids from IFNα14 for the treatment of HBV infection. Again, differentiated HepaRG cells were infected with HBV and stimulated with different concentrations of the IFNα2-mutants, which were mutated in their binding region to IFNAR1 (IFNα2-EIFK) or to IFNAR2 (IFNα2-PF) or both (IFNα2-PEIFKF), the putative tunable anchor (IFNα2-QASMT), or outside of these regions (IFNα2-NNEMM). Stimulation with IFNα14 significantly reduced HBsAg levels (IC_50_[U/ml]: 34.1) in contrast to the low activity of IFNα2 (IC_50_[U/ml]: 514.1) (Fig. 2C, D). Similar to previous results, we measured the highest antiviral activity with IFNα2-EIFK (IC_50_[U/ml]: 8.2) in which four amino acids were changed that are required for optimal binding to IFNAR1 (9). Interestingly, site-directed mutagenesis of the region outside of the receptor binding sites and the putative tunable anchor region also improved the anti-HBV activity of IFNα2 (IFNα2-NNEMM; IC_50_[U/ml]: 59.8), whereas modification in the putative tunable anchor region alone had no effect on the antiviral activity of IFNα2 (IFNα2-QASMT; IC_50_[U/ml]: 96.19).

### Combined mutations in IFNAR1 binding sites, the putative tunable anchor region, and outside these regions influenced the anti-HIV activity of IFNα2

We previously reported a significantly higher anti-HIV potency of IFNα14 and IFNα6 in comparison to IFNα2 (11, 20). Thus, we determined the impact of critical IFN residues on HIV replication in TZM-bl cells.

Titration experiments with parental IFNα2 and IFNα6 showed a stronger antiviral effect of IFNα6 (IC_50_[U/ml]: 64.99) than IFNα2 (IC_50_[U/ml]: 2001) (Fig. 3A, B). Next, we used the IFNα2-mutants IFNα2-MS (mutated in the IFNAR2 binding sites) and IFNα2-ASVT (mutated in the putative TA region) in the same assay. Only targeted modifications of the putative tunable anchor region of IFNα2 (IFNα2-ASVT) slightly improved the antiviral activity of IFNα2 against HIV (IC_50_[U/ml]: 363.4), but this was not statistically significant. Insertion of the IFNα6 residues which are important for IFNAR2 binding had no effect on the inhibition of HIV replication compared to parental IFNα2.

**Figure 3:**
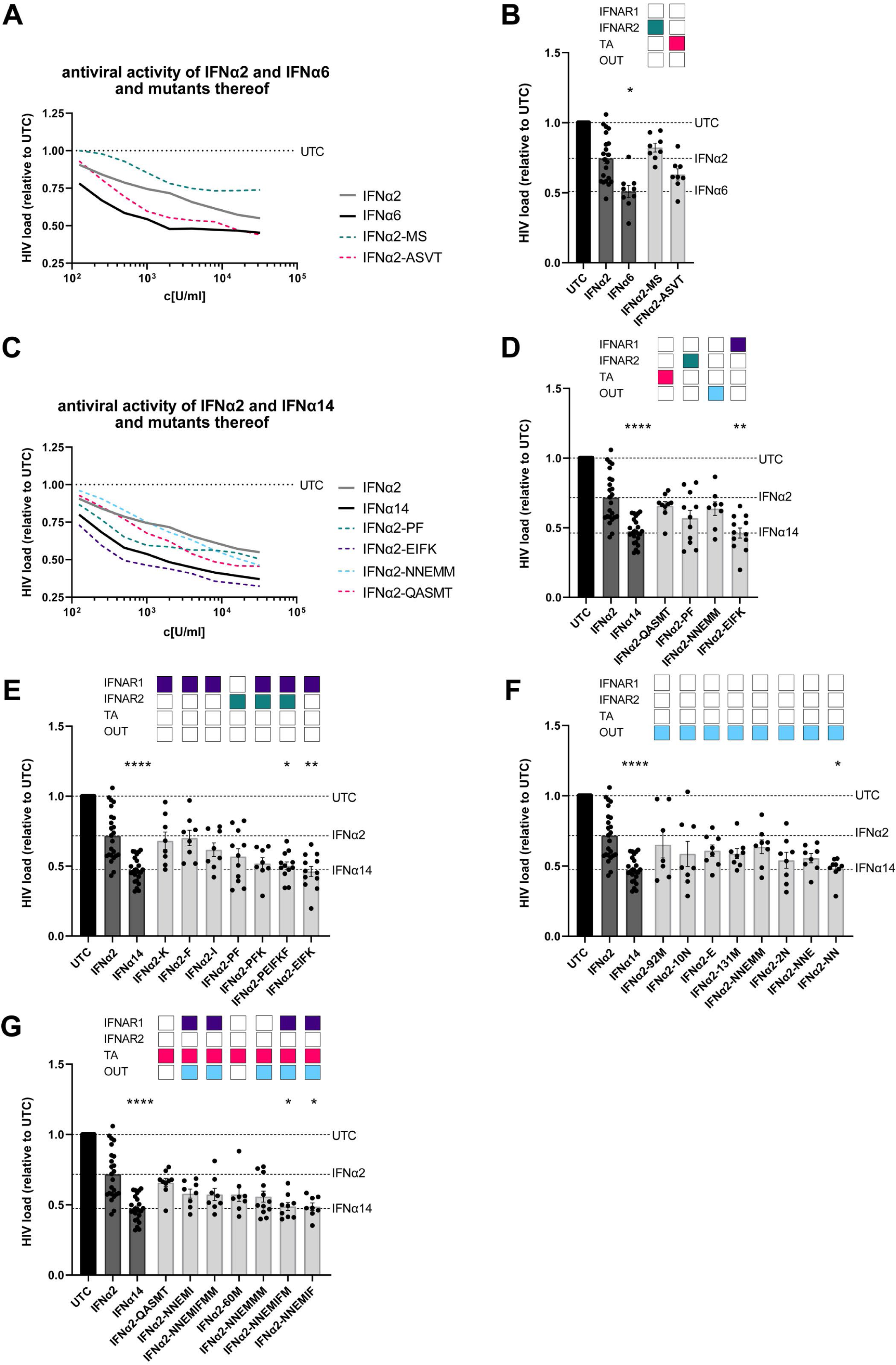
The antiviral activity of IFNα2-mutants against HIV. TZM-bl cells were infected with a R5-HIV-1_NL4-3-IRES-Ren_ reporter virus at a MOI of 0.02 and treated with IFNα2, IFNα6, IFNα14, or IFNα2-mutants. (A, C) IFNα2, IFNα6, IFNα14, and IFNα2-mutants were serially diluted starting at a concentration of 32,000 U/ml to measure dose-dependent anti-HIV effects at 3 days post infection. (B, D-G) TZM-bl cells were treated with 2,000 U/ml of the designated IFNs and IFNα2-mutants to screen for their anti-HIV activity by quantifying luciferase activities. (A,C) n = 6; (B, D-G) n = 8-20; Mean ± SEM is plotted. Kruskal-Wallis test was performed against untreated control (UTC). Statistical significance is depicted as * p < 0.05, **p < 0.001 ****p < 0.0001. TA = Putative tunable anchor; OUT = outside

Next, we evaluated potential critical residues in IFNα14, to improve the anti-HIV activity of IFNα2. As depicted in Fig. 3C and D, stimulation with the IFNα2- mutants, which are mutated in their binding region to IFNAR1 (IFNα2-EIFK), to IFNAR2 (IFNα2-PF), the putative tunable anchor (IFNα2-QASMT), or outside of these regions (IFNα2-NNEMM) showed differential antiviral activities. Targeted mutations of the whole IFNAR1 binding site (D82E; T86I; Y89F; R120K) of IFNα2 significantly improved the antiviral activity against HIV, whereas mutations outside the three described regions or at the IFNAR2 binding site alone had only minor, non-significant effects on the suppression of HIV replication (Fig. 3C, D). We further analyzed single or multiple mutations in the IFNAR1/2 binding region (Fig. 3E), outside of the described protein motifs (Fig. 3F), and combinations of mutations targeting IFNAR1, the putative tunable anchor, and the outside regions together (Fig. 3G). Single mutations at the IFNAR1 binding sites (IFNα2-K; IFNα2-F; IFNα2-I) did not enhance the anti-HIV activity of IFNα2 (Fig. 3E). Only the mutation of all IFNAR1 binding sites (D82E; T86I; Y89F; R120K) and the combination with the two IFNAR2 binding sites (L26P; L153F) resulted in a significant reduction in HIV replication (Fig. 3E), which was also confirmed by lower IC_50_ values compared to the parental IFNα2 (IC_50_[U/ml]: 88.73 (IFNα2- EIFK); 41.56 (IFNα2-PEIFKF); 761.5 (IFNα2); Supp. Fig. 2). Of note, the residues 2 (D) and 10 (G) might also be important for the regulation of the anti- HIV activity of IFNα2, as the combined amino acid changes D2N and G10N outside of the defined regions of interest significantly reduced HIV loads, whereas stimulation with parental IFNα2 had no significant effect on HIV loads measured by Renilla luciferase activity (Fig. 3F, Supp. Fig. 2). We also analyzed combinations of mutated residues critical for IFNAR1 binding, the putative tunable anchor and outside regions. Changing at least six amino acids in IFNα2 to amino acids from IFNα14 (D2N; G10N; G37E; I60M; T86I; Y89F) completely converted the antiviral activity of IFNα2 to the much stronger IFNα14 activity (Fig. 3G, Supp. Fig. 2; IFNα2-NNEMIF(M)). Our data suggest that for the antiviral activity of IFNα against HIV, the residues critical for binding to IFNAR1 are the most important. In addition, also the two residues close to the N-terminus of IFNα2 (Asp 2; Gly 10), which are outside the three defined regions of interest, strongly influence the anti-HIV activity of IFNα.

### Combined mutations in the IFNAR1 binding sites, putative tunable anchor region, and outside of these defined regions are required for potent activation of downstream signaling cascades in PBMCs

To elucidate their antiviral effects and molecular signaling pathways in primary cells, IFNα2 mutants, which significantly reduced HIV loads in infected TZM-bl cells (IFNα2-PEIFKF; IFNα2-EIFK; IFNα2-NN; IFNα2-NNEMIF; IFNα2-NNEMIFM), were further analyzed in primary HIV target cells. In order to further scrutinize the biological effects of type I IFNs during HIV infection, we utilized PBMCs from healthy individuals and infected these cells *in vitro* with a Renilla Luciferase expressing X4- or R5-tropic HIV_NL4-3_ which resulted in comparable infection levels (data not shown). Firstly, titrations with parental IFNα2 and IFNα14 were performed to determine the appropriate concentrations. As depicted in Figure 4A, a dose-dependent inhibition of HIV was evident, with IFNα14 exhibiting stronger inhibition compared to IFNα2. Next, HIV-infected cell cultures were directly stimulated with two different concentrations (325U/ml and 2000U/ml) of parental IFNα2, IFNα14, or IFNα2-mutants and the IFN-mediated effects on viral loads were analyzed 3 days post infection (dpi) (Fig. 4B-E). According to the infections of TZM-bl cells (Fig. 3), IFNα2-EIFK, in which IFNAR1 binding sites were introduced from IFNα14, as well as the combination of mutated residues at the IFNAR1 binding sites, putative tunable anchor region, and mutations outside of these defined motifs significantly reduced HIV infection in PBMCs compared to IFNα2 (Fig. 4C, E). Both the parental IFNs and the IFNα2-mutants exhibited similar antiviral efficacy in X4- and R5-tropic virus infections (Fig. 4B-E).

**Figure 4:**
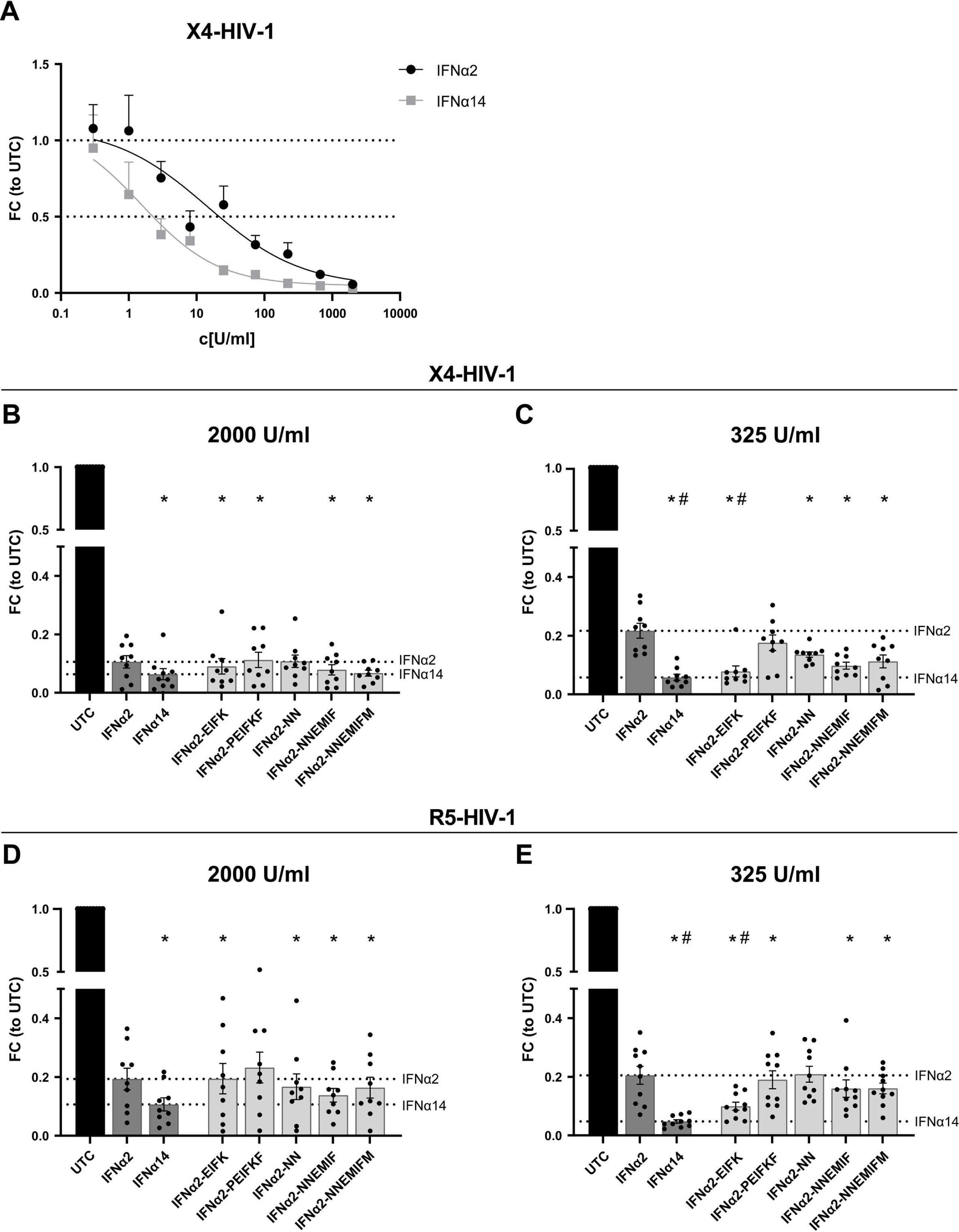
The anti-HIV activity of IFNα2-mutants in primary cells. (A) PBMCs were infected with X4-HIV-1_NL4-3-IRES-Ren_ at a MOI of 0.25. IFNα2 and IFNα14 were titrated starting at a concentration of 2,000 U/ml in a 3-fold dilution. (B, D) PBMCs were infected with a X4- or R5-tropic HIV-1_NL4-3-IRES-Ren_ reporter virus at a MOI of 0.25 and treated with 2,000 U/ml or (C, E) 325 U/ml of the designated IFNs and IFNα2-mutants. To evaluate the anti-HIV capacity the luciferase activity was measured 3 days post infection. Mean values ± SEM are shown for n=9-10. Statistical analyses between the different groups were done by using non-parametric Friedman test and Dunn’s multiple comparison test. Statistical significance is depicted as * p < 0.05 against untreated control (UTC) and # p < 0.05 against parental IFNα2.

As binding of type I IFNs to their receptors leads to the induction of multiple signaling pathways (17, 23), we performed phosphoflow analysis to elucidate the underlying molecular mechanism of the different effector responses after IFNα treatment of HIV-infected PBMCs. To this end, we stimulated PBMCs from healthy individuals with the different parental IFNα subtypes and IFNα2-mutants for 15 min and analyzed the phosphorylation of the immune cell signaling molecules STAT1, STAT3, and STAT5 in CD4^+^ and CD8^+^ T cells. Treatment with IFNα14 strongly increased the frequencies of phosphorylated STAT1^+^ T cells, which increased only slightly after stimulation with IFNα2. All IFNα2-mutants increased the frequencies of pSTAT1^+^ CD4^+^ and CD8^+^ T cells compared to IFNα2; however, significance was only reached for the IFNα2-NNEMIFM molecule (Fig. 5A, D). For STAT3 phosphorylation in T cells, we solely observed a significant increase in percentages for IFNα14- and IFNα2-PEIFKF-treated CD8^+^ T cells compared to untreated controls (Fig. 5B, E), indicating that certain IFNα residues influence the activation of specific downstream signaling pathways. We also determined the frequencies of pSTAT5^+^ T cells after stimulation with the different IFNs. Similar to the results observed for pSTAT1 (Fig. 5A, D), treatment with IFNα14 resulted in significantly increased percentages of pSTAT5^+^ T cells, which was only slightly influenced by the parental IFNα2 (Fig. 5C, F). Again, stimulation with IFNα2-PEIFKF and IFNα2- NNEMIFM significantly enhanced the frequencies of pSTAT5-expressing T cells comparable to the results with IFNα14. These data nicely demonstrate that the combined modulation of the IFNAR1/2 binding sites (IFNα2-PEIFKF) or the IFNAR1 binding site and the putative tunable anchor and motifs outside these regions (IFNα2-NNEMIFM) resulted in an augmented activation of important downstream signaling cascades, even those that are distinct from the canonical type I IFN signaling pathway STAT1/STAT2. This further highlights the importance of certain amino acid motifs for the different biological activities of the IFNα subtypes. Such critical residues may be responsible for the qualitative differences of the numerous IFNα subtypes.

**Figure 5:**
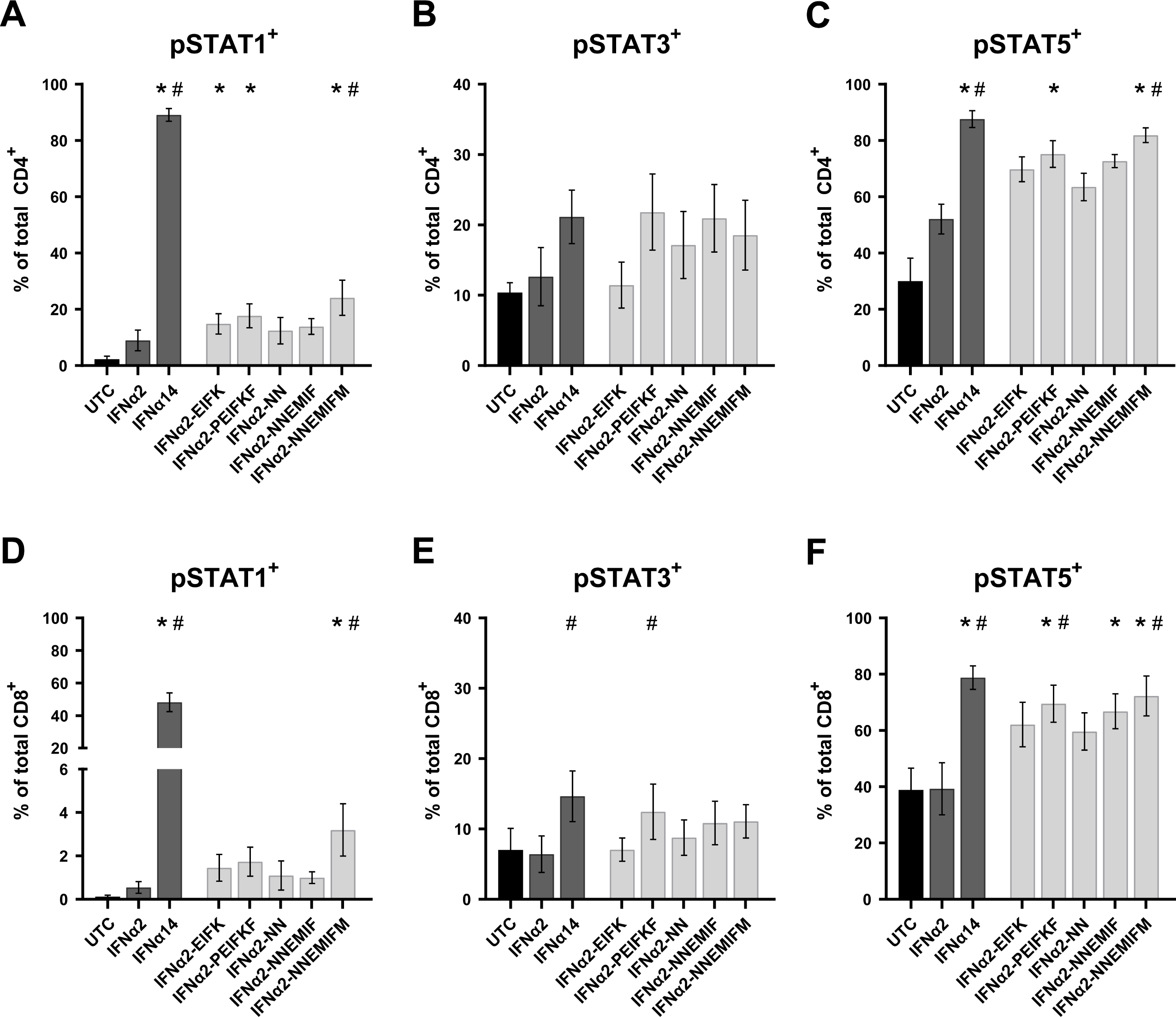
Potency of IFNα2-mutants to activate STAT molecules. PBMCs were stimulated with 2,000 U/ml of the designated IFNs and IFNα2- mutants or without IFN (UTC) for 15 min in presence of the surface markers anti- CD3, anti-CD4, anti-CD8, and the viability marker FVD. Cells were then fixed and permeabilized for phosphostaining with anti-STAT1 pTyr701 (A, D), anti-STAT3 pTyr705 (B, E), and anti-STAT5 pTyr694 (C, F). Mean values ± SEM are shown for n=6. Statistical analyses between the different groups were done by using non-parametric Friedman test and Dunn’s multiple comparison test. Statistical significance is depicted as * p < 0.05 against untreated control (UTC) and # p < 0.05 against parental IFNα2.

## Discussion

Type I IFNs play a pivotal role in the host defense against infectious agents. In humans, the type I IFN family comprises IFNβ, IFNε, IFNκ, IFNω, and twelve IFNα subtypes. The overall homology of the IFNα proteins ranges from 75 to 95% amino acid sequence identity. Despite binding to the same cellular receptor, consisting of the two subunits IFNAR1 and IFNAR2, their antiviral, immunomodulatory, and antiproliferative potencies differ considerably (6, 9–11). It is largely elusive why different IFNα proteins exhibit distinct effector functions. Different receptor affinities and/or interaction interfaces within the receptor have been discussed, which may account for the observed variability in the biological activity (24, 25). Furthermore, dosage, cell type, timing, and the present cytokine milieu might further affect the IFN effector response (15). In particular, the antiviral activity of clinically approved IFNα2 versus other IFNs like IFNα14 or IFNα6 is rather low, which might explain the overall moderate outcome of IFNα2- therapy in chronic viral infections and the occurrence of severe side effects. Thus, new and safe IFN therapeutics with broad or target-specific antiviral activity have to be discovered.

Here, we focused on the structural differences between the highly antiviral IFNα subtypes IFNα6 and IFNα14, and the clinically approved IFNα2. IFNα6 expresses 21 different amino acids, compared to IFNα2b (Supp. Fig. 1), and only two of these different residues are located in regions that are critical for receptor binding (IFNAR2). Although IFNAR1 binding regions (13) are identical between IFNα2b and IFNα6, their affinities to IFNAR1 differ strongly (3.8 µM for IFNα2b and 0.83 µM for IFNα6) (24), suggesting that other residues might also influence receptor binding. In contrast, the sequence of IFNα14 harbors 28 variant amino acids compared to IFNα2b with four differences in the IFNAR1 binding region (positions 82, 86, 89 and 120; IFNα2 numbering) and 2 differences in the IFNAR2 binding region (13, 26–28) (positions 26 and 153; IFNα2 numbering). These amino acid variations result in stronger binding affinity of IFNα14 to IFNAR1 (0.68 µM) and IFNAR2 (0.7 nM) compared to IFNα2b (3.8 µM and 1.3 nM, respectively) (24). Our structural analysis further yielded a region in helix B (T52-L66) within the IFNα protein with a surprising conservation pattern, mainly conserved residues at the outside facing IFNAR1 but more variable at the inside facing the core of IFNα. This could serve as a putative “tunable anchor”, i.e., while the binding site with IFNAR1 is conserved, its position and fine structure might be modified by targeted mutations in the core facing region. In particular, antiviral activity against HIV, as well as phosphorylation of STAT1 and STAT5 (Fig. 4 and 5) significantly improved when a mutation (I60M) within the newly identified putative TA region in combination with a single mutation in the IFNAR1 binding region and mutations outside the regions of interest were inserted. Another IFNα2-mutant (IFNα2-YNS) was previously identified by phage display with very tight binding affinity against IFNAR1 (29). The YNS mutations (H57Y, E58N, Q61S) were inserted at the conserved IFNAR1 binding site, which is in direct proximity of the more variable residues of the putative tunable anchor. A 60-fold increase in IFNAR1 binding and thus a 150-fold higher antiproliferative activity in human WISH cells as well as a strong anti-tumor effect *in vivo* was observed. The antiviral activity against Vesicular stomatitis virus was only slightly increased, i.e., in particular the antiproliferative activity of IFNs correlates with IFNAR1 binding affinity. Similar results were reported with the IFNα2-mutant HEQ, where the positions mentioned above were mutated to alanine, resulting in an increased IFNAR1 binding affinity comparable to IFNβ and a corresponding enhanced antiproliferative activity, but only slightly increased antiviral potency (30). These data suggest that, especially, the IFNAR1 binding region within the helix T52-L66 strongly affects the IFN-response. Thus, a detailed analysis of the putative TA residues and their impact on antiproliferative and immunomodulatory effects is indispensable. In particular, the influence of the putative TA region on IFNAR1 and IFNAR2 binding affinities is mandatory to fully understand the biological role of this region within the IFN core. Of note, also other regions are critical for IFNAR1 binding, which have to be considered as regulators for the differences in biological activity between the different IFNα subtypes. Position 120, which belongs to the IFNAR1 binding region, was reported to be critical for the antiviral activity. This position is conserved in most subtypes (Arg 120), but IFNα1/13, IFNα14 and IFNα21 express Lys at that position and site-directed mutagenesis of R120K in IFNα2 or IFNα4 significantly increased the antiviral activity against Semliki Forest virus (31–33). In addition, amino acid region 81– 95, which harbors multiple critical residues for IFNAR1 binding, was shown to be important for antiproliferative activity (34). Although IFNα6 and IFNα14 exhibit a comparable strong antiviral activity against HIV-1 and HBV, their amino acid sequence, especially at the important IFNAR1 binding site, differs strongly.

Specifically, IFNα6 possesses identical amino acids at the IFNAR1 binding site as IFNα2, yet it exhibits significantly stronger binding affinity to IFNAR1 (24) and displays much higher antiviral activity than IFNα2 (as shown in Fig. 2A, B and Fig. 3A, B). One possible explanation could be the minimal unique structural characteristics of each IFNα subtype, which may lead to distinct interactions with IFNAR1. These interactions could be influenced by the three-dimensional arrangement of critical amino acids within the binding site or the putative TA, causing variations in the strength of binding and subsequent downstream signaling pathways.

Further, we could show that different signaling cascades, including the classical STAT1, as well as non-classical STAT3 and STAT5, were strongly phosphorylated by IFNα14 in T cells. The combined modulation of the IFNAR1/2 binding sites (IFNα2-PEIFKF) or the IFNAR1 binding site and the putative TA and motifs outside these regions (IFNα2-NNEMIFM) resulted in an increased activation of important downstream signaling cascades in T cells (Fig. 5). The p- STAT response does not show significant differences among the various IFN mutants, despite variations in their antiviral responses. One possible explanation could be that the differences in antiviral activity between the IFN mutants are not solely mediated by the classical JAK-STAT signaling pathway. Other signaling pathways or alternative mechanisms may be involved in modulating the antiviral response, which are not directly reflected in the p-STAT1/3 or 5 levels. Additionally, the antiviral activity of IFNs can also be influenced by factors beyond the JAK-STAT pathway, such as the distinct pattern of induced ISGs or the activation of other downstream effectors. These alternative pathways may contribute to the observed differences in antiviral responses among the IFN mutants. The IFNα2-YNS mutant was previously analyzed for its potency to induce phosphorylation of STAT1, STAT3, and STAT5 molecules in PBMCs, as well as for its antiviral and antiproliferative activity (13). Again, the increase in antiviral activity of IFNα2-YNS compared to IFNα2 was low in contrast to the anti- proliferative effect, which was up to 1000-fold increased. Although the potency to phosphorylate the different STAT molecules was comparable between the different IFNs (IFNα2, IFNα2-YNS, IFNα7 and IFNω), the IFNα2-YNS mutant had a lower EC_50_ for STAT1 phosphorylation, implying that stronger binding to IFNAR1 leads to preferential activation of the classical JAK-STAT pathway. However, the influence of the YNS mutations within IFNα2 on the antiviral activity was rather low and comparable to the effects on STAT phosphorylation. This further strengthens the hypothesis that modulation of the IFNAR1 binding region mainly impacts the so-called tunable activity of IFNs that are defined as antiproliferative and immunomodulatory activities of IFNs (35). The tunable activity may vary between different cell types and is often stronger induced by high-affinity binders and most likely requires higher IFN concentrations, longer times of IFN stimulation, and higher receptor surface concentrations(35, 36). In contrast, the antiviral activity of IFNα is defined as robust activity stimulated by low amounts of IFNα, low receptor surface expression, and a common program in all cells mainly mediated by the classical JAK-STAT-driven ISRE gene transcription. For both viral infections (HIV, HBV), we observed the highest improvements in antiviral activity when all four IFNAR1 binding sites (positions 82, 86, 89 and 120), which differ between IFNα2 and IFNα14, were mutated. The resulting mutant IFNα2-EIFK exhibited an antiviral potency comparable to IFNα14 (Fig. 2, 4), which was not further improved by modulating the IFNAR2 binding sites. This supports data describing another IFNα2-YNS mutant with an increased binding affinity to IFNAR2 by exchange of its C-terminal tail (IFNα2- YNS-α8tail) (37). This exchange further boosted the antiproliferative activity, whereas the robust antiviral activity was not affected at all. For the antiviral activity of IFNα subtypes many residues (positions 30, 33, 77, 78, 123, 129, 130, 133, 134, 135, 140; IFNα2 numbering) were reported to be critical (38); however, these residues are all conserved in all IFNα subtypes. Up to now, the exact mechanism that regulates binding to IFNAR2 is not completely understood. Four conserved residues appear to be essential for the interaction with IFNAR2 (positions 30, 33, 148, and 149; IFNα2 numbering), but it seems that small variations in close proximity to these positions modulate the IFNAR2 binding affinity (39).

Here, combining IFNα sequence variability and 3D structural data, we suggest how IFNα variants could achieve differential functions. While the overall binding can be fixed by a set of conserved anchors, other positions are more variable, allowing for a tuning of receptor affinities, or of on-/off-rates for complex formation of IFNα and receptor molecules. These data give rise to the development of new therapeutic IFN molecules with high antiviral activity against a variety of different viruses.

## Materials and Methods

### Sequence and structural analysis of IFNs

Sequences of human IFNα1 (NP_076918.1), IFNα2 (NP_000596.2), IFNα4 (NP_066546.1), IFNα5 (NP_002160.1), IFNα6 (NP_066282.1), IFNα7 (NP_066401.2), IFNα8 (NP_002161.2), IFNα10 (NP_002162.1), IFNα14 (NP_002163.2), IFNα16 (NP_002164.1), IFNα17 (NP_067091.1), and IFNα21 (NP_002166.2) were aligned with MAFFT (22) using the FFT-NS-2 strategy.

To map sequence entropies on IFNα2 structure (PDB entry 1itf, (19)), the relevant parts of the above IFNα sequence alignment (including the signal peptide), namely positions 24-66 and 68-189 were firstly extracted with 1 Emboss program extractalign, version 6.6.0.0 (40). Sequence entropies were computed for all positions of the extracted alignment with function entropy of R-package bio3d, version 2.4.0 (41) using the standard entropy *H*_*j*_ at alignment position

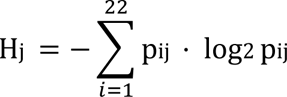

with relative frequencies *pij* of amino acid symbols *i* at position *j*, and *i* = 1, 2, …, 22 for the 1-letter amino acid codes and additionally a letter for nonstandard amino acids and a gap symbol. Entropies were mapped on IFNα2 structure (PDB entry 1itf; NMR model 1, (19)), with R-script (42). Structure alignments and RMSD calculations were performed with pymol 1.8 (43).

### Site-directed mutagenesis and IFN production

Site-directed mutations were introduced using the primers listed in Tab. 2 and the QuikChangeII kit (Agilent) according to manufacturer’s instructions. All constructs were confirmed by sequence determination of the coding sequence.

**Table 2:**
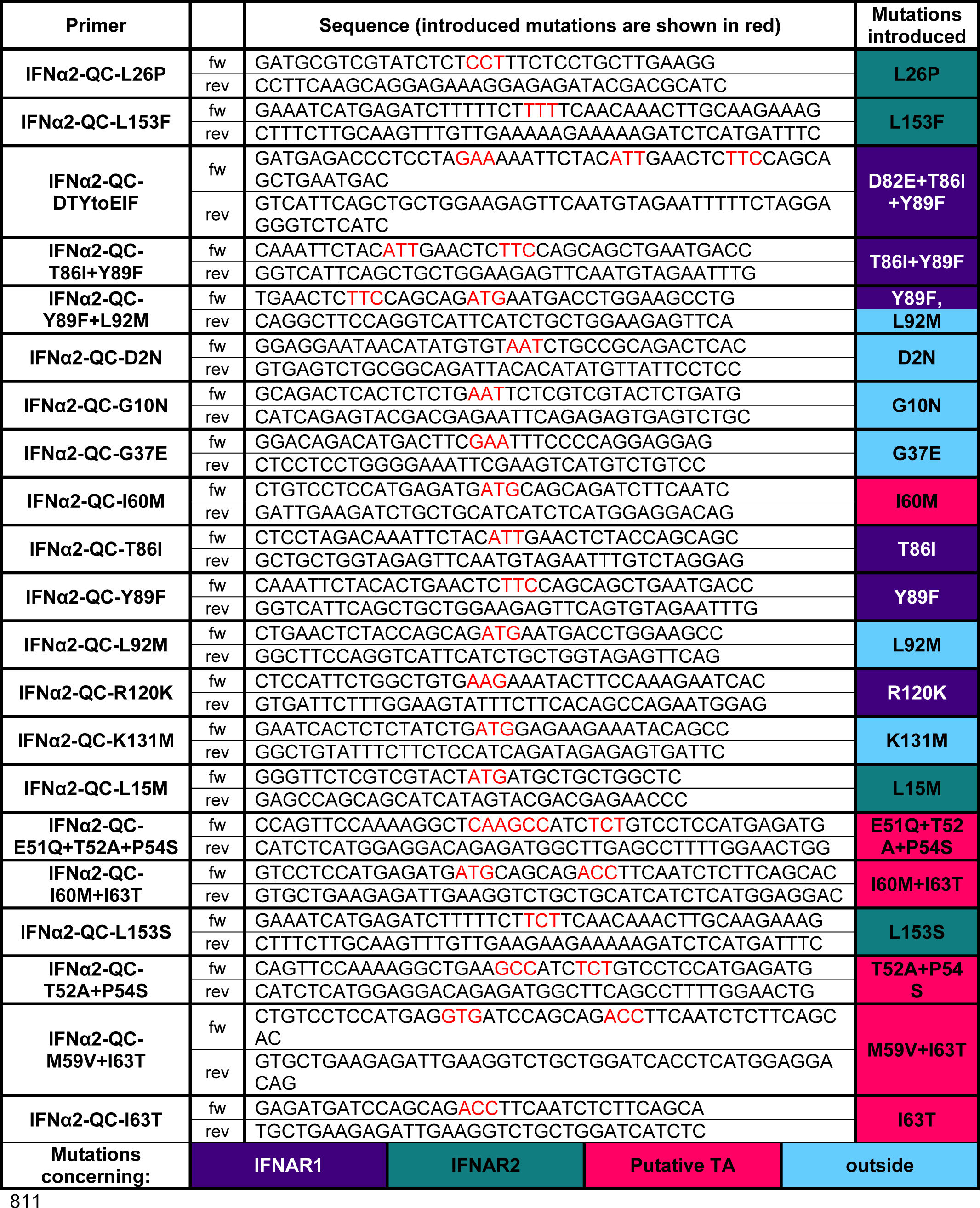
Used Primers.

Human IFNα subtype and mutant genes were optimized for expression in *Escherichia coli*. Isolated inclusion body proteins denatured with guanidine hydrochloride were refolded in arginine refolding buffer and purified by anion- exchange and size exclusion chromatography (29). Protein concentrations were determined using Nano-Drop 2000c (Thermo Scientific, Wilmington, DE), and endotoxin levels were less than 0.0025 endotoxin units (EU)/ml (ToxinSensor; Genscript, Piscataway, NJ). These laboratory-produced proteins were used for all experiments.

Because the standard biological method to quantify interferons is with antiviral assays, we were concerned that the differential antiviral effects of the various interferon subtypes might produce aberrant results. Therefore, ISRE-luc reporter cells (11) were grown for 24 h before adding serial dilutions of our recombinant IFNα subtypes and commercially available IFNα subtypes (PBL Assay Science) for 4.5 h. The cells were lysed with lysis buffer (pjk) and Firefly luciferase activity was measured subsequently. Six experiments were done comparing the stated activities of commercially available IFNα subtypes (PBL Assay Science, Piscataway, NJ) with relative light units (RLU) obtained from our ISRE assay. All the units given in the text correspond to PBL units. PBL determines the activities of interferons using a cytopathic inhibition assay on the human lung carcinoma cell line A549 with encephalomyocarditis virus (EMCV).

### Isolation and cultivation of primary cells

HIV-negative blood samples (n=6-9) were donated by healthy individuals of the University Hospital Essen. Blood collection was approved by the Ethics Committee (No.:11-4715) of the University of Duisburg-Essen.

PBMCs were isolated from each blood sample by density gradient centrifugation as described elsewhere (7).

### Infection with HIV

X4-HIV-1_NL4-3-IRES-Ren_ and R5-HIV-1_NL4-3-IRES-Ren_ reporter viruses were produced by transfection of HEK293T cells with pNL4-3Ren. The TCID_50_ were calculated by X-Gal staining of infected TZM-bl reporter cells.

PBMCs were cultivated at a density of 1x10^6^ cells/ml in RPMI 1640 supplemented with 10% FCS, 100 U/ml penicillin and 100 µg/ml streptomycin, 2 mM L-gultamine, and 10 mM HEPES. Additionally, PBMCs were activated with 1 µg/ml PHA in presence of 10 ng/ml IL-2 (Miltenyi Biotec). PBMCs were mock- treated or infected with multiplicity of infection (MOI) of 0.25 via spinoculation at 1,200 x g for 2 h. Viral input was removed and cells were washed with PBS. Cells were cultivated in fresh media containing 2,000 U/ml or 325 U/ml of the appropriate IFNα subtypes or IFNα2-mutants at a density of 1x10^6^ cells/ml. Additionally, IFNα2 and IFNα14 were titrated on X4-HIV-1_NL4-3-IRES-Ren_ infected PBMCs starting at a concentration of 2,000 U/ml in a 3-fold dilution. Cells were lysed 3 dpi with lysis buffer (pjk) for the determination of viral loads.

### TZM-bl Assay

TZM-bl cells were seeded at a density of 5,000 cells per well in a 96-well plate in DMEM containing 10% FCS, L-glutamine, 100 U/ml penicillin, and 100 µg/ml streptomycin. Next day, cells were infected with a R5-HIV-1_NL4-3-IRES-Ren_ reporter virus with a MOI of 0.02 and treated with 2,000 U/ml of the appropriate IFNα subtypes or IFNα2-mutants for 72 h. Additionally, the indicated IFNs were titrated on TZM-bl cells starting at a concentration of 32,000 U/ml in a 2-fold dilution. The infection efficacy was then analyzed by a luciferase assay according to the manufacturers standard protocol (pjk Renilla-Juice Luciferase Assay).

### Infection with HBV

Fully differentiated HepaRG cells (44) were infected with HBV genotype D at an MOI of 500 genome equivalents (geq) in inoculation media (differentiation media, 10% PEG8000), followed by 20 h incubation. Subsequently, the cells were washed twice with PBS and 300 µl of differentiation media were added. Cells were stimulated on days 0, 1, 4 and 6 post-infection. Supernatant was collected on day 9 post-infection.

### Hepatitis B surface antigen ELISA

The supernatants were analyzed using the Hepatitis B surface antigen Ab ELISA Kit (Abnova) according to the manufacturer’s instructions. Serially diluted positive control (8 ng/ml) was used to generate a standard curve. The absorbance was measured at 450 nm with a reference wavelength of 620 nm using Spark® 10M multimode microplate reader (Tecan).

A standard curve was generated using the Four Parameter Logistic (4PL) Curve Calculator (AAT bioquest). The 4PL curve was used to calculate the amount of HBs in the supernatants.

### Phosphoflow analysis

For phosphoflow analysis cells were stimulated with 2,000 U/ml IFNα2, IFNα14, IFNα2-mutants, or left unstimulated (UTC) for 15 min at 37°C. Surface staining was performed simultaneously to IFNα stimulation with the following antibodies: anti-CD3 (UCHT1, eBioscience™), anti-CD4 (RPA-T4, BioLegend), anti-CD8 (RPA-T8, BioLegend), as well as FVD for exclusion of dead cells. After stimulation cells were immediately fixated with pre-warmed Fixation Buffer (BioLegend) at 37°C. Cells were then permeabilized with pre-chilled TruePhos™ Perm Buffer (BioLegend) at -20°C for 1 h. Subsequently, cells were washed twice with FACS Intracellular Staining Perm Wash buffer and the following antibodies were added to the cells: anti-STAT1 pTyr701 (Miltenyi Biotec), and anti-STAT3 pTyr705 (eBioscience™), and anti-STAT5 pTyr694 (BioLegend). After a 30 min incubation, cells were washed twice with FACS Intracellular Staining Perm Wash buffer and stored at 4°C until acquisition. Samples were acquired with a BD LSR II flow cytometer with a HTS module and data were analyzed using FACSDiva and FlowJo Version 10.8.

### Statistical analysis

Experimental data were reported as means ±SEM. Statistically significant differences between treated and untreated or IFNα2 treated primary cells were analyzed using non-parametric Friedman test and Dunn’s multiple comparison test. Kruskal–Wallis one-way analysis of variance on ranks with Dunn’s multiple comparison procedure was used to monitor statistical differences in cell culture conditions. Dose-response analyses were performed to evaluate inhibitory concentrations (IC_50_). All analyses were performed using GraphPad Prism software v8 (GraphPad, San Diego, CA, USA).

## Acknowledgments

This work was supported by the DFG to K.S. (SU1030/2-1; SU1030/1-2) and U.D. (DI714/18-2). We are grateful to the Stiftung Universitätsmedizin Essen of University Hospital Essen for financial support of this study. This project was supported by the Sino-German Virtual Institute for Viral Immunology (SGVIVI). We acknowledge support by the Open Access Publication Fund of the University of Duisburg-Essen.

